# Reevaluating claims of ecological speciation in *Halichoeres bivittatus*

**DOI:** 10.1101/2021.03.11.435027

**Authors:** Dan L. Warren, Ron I. Eytan, Alex Dornburg, Teresa L. Iglesias, Matthew C. Brandley, Peter C. Wainwright

**Affiliations:** Biodiversity and Biocomplexity Unit, Okinawa Institute of Science and Technology Graduate University, Okinawa, Japan..; Department of Marine Biology, Texas A&M University at Galveston, Galveston, TX, USA..; Department of Bioinformatics and Genomics, University of North Carolina Charlotte, Charlotte, NC, USA..; Animal Resources Section, Okinawa Institute of Science and Technology Graduate University, Okinawa, Japan..; Carnegie Museum of Natural History, Pittsburgh, PA, USA..; Department of Evolution and Ecology, University of California, Davis, CA, USA..

## Abstract

Understanding the role of ecological processes in speciation has become one of the most active areas of research in marine population biology in recent decades. The traditional view was that allopatry was the primary driver of speciation in marine taxa, but the geography of the marine environment and the dispersal capabilities of many marine organisms render this view somewhat questionable. One of the earliest and most highly cited empirical examples of ecological speciation with gene flow in marine fishes is that of the slippery dick wrasse, *Halichoeres bivittatus*. Evidence for this cryptic or incipient speciation event was primarily in the form of a deep north-south divergence in a single mitochondrial locus, combined with a finding that these two haplotypes were associated with different habitat types in the Florida Keys and Bermuda, where they overlap. Here we examine habitat assortment in the Florida Keys using a broader sampling of populations and habitat types than were available for the original study, and find no evidence to support the claim that haplotype frequencies differ between habitat types, and little evidence to support any differences between populations. These results severely undermine claims of ecological speciation with gene flow in *Halichoeres bivittatus*. We argue that future claims of this type should be supported by multiple lines of evidence that illuminate potential mechanisms and allow researchers to rule out alternative explanations for spatial patterns of genetic differences.

## Introduction

It is widely recognized that many marine organisms challenge traditional models and expectations of the geography of speciation. Marine environments are host to a great deal of the world’s biodiversity, yet the relative rarity of obvious barriers to dispersal coupled with long pelagic larval durations offer few opportunities for long-term genetic isolation or population structuring that are generally thought to be prerequisites of the speciation process (Palumbi 1994, 1992). Not surprisingly, understanding the processes leading to speciation and the accumulation of biodiversity in marine environments has been an area of intense research in recent decades (Cowman et al. 2017; Hodge and Bellwood 2016; Bowen et al. 2013; Gaboriau et al. 2018; Faria, Johannesson, and Stankowski 2021). Some have argued that speciation in marine systems is primarily allopatric –as in terrestrial systems (Mayr 1954)– with barriers to gene flow being more cryptic in the ocean than on land (Taylor and Hellberg 2006; Goetze 2011, 2005). In contrast, a number of recent case studies have suggested that ecological processes likely play a major role in promoting speciation in marine systems, often with gene flow (Rocha et al. 2005; Prada and Hellberg 2020; Taylor and Hellberg 2005; Whitney, Donahue, and Karl 2018; Momigliano et al. 2017; Teske et al. 2019; Faria, Johannesson, and Stankowski 2021).

In a pioneering study, Rocha et al. (Rocha et al. 2005) presented evidence supporting the possibility of ecological speciation in coral reef fishes, presenting two possible cases of parapatric speciation in Atlantic *Halichoeres*. One of these case studies focused on *Halichoeres bivittatus*, in which they demonstrate a deep divergence in cytochrome B sequences between a northern lineage (spanning the northern Gulf of Mexico, peninsular Florida, and the eastern coast of the United States), and a southern lineage (spanning the Yucatan peninsula, Cuba, the eastern Bahamas and all points south including the southern Caribbean and coastal Brazil). Finding a deep divergence at a locus with geographic structure is not in itself evidence of speciation. For example, such a divergence can be expected even under neutral processes (Irwin 2002), in particular with respect to mitochondrial loci such as cytochrome B (Irwin 2002; Taylor and Hellberg 2006; Neigel and Avise 1993). However, Rocha et al. (2005) presented evidence that the two haplotypes were preferentially associated with different types of habitat in the Florida Keys and Bermuda, and presented several lines of evidence arguing that there was significant potential for gene flow between northern and southern populations. This finding of divergence in the face of gene flow combined with habitat partitioning in the contact zone between the two haplotypes led the authors to conclude that ecological processes either had driven, or were in the process of driving, parapatric speciation in this system. If true, this represents a departure from the more common pattern of speciation in this clade (Wainwright et al. 2018) and other Caribbean fishes, in which new species seem to primarily arise from vicariance or long distance dispersal events (Robertson et al. 2006; Choat et al. 2012). However, support for habitat segregation in the Florida Keys was based on samples of only two populations and included larval samples, which may demonstrate patterns of spatial segregation due to reasons that are not informative for questions of speciation. Here we present the results of an attempt to further explore patterns of habitat segregation for *H. bivittatus* in the Florida Keys by sampling additional populations of adults and conducting more extensive statistical analyses.

## Methods

To test habitat partitioning among *Halichoeres bivittatus* haplotypes, we analyzed the same mitochondrial cytochrome B fragment as Rocha et al. (2005) for thirteen additional populations/collection sites. We sampled eight populations in the Florida Keys including four populations on the edge of the continental shelf (Sombrero Light, 11 Foot Mound, XMuta, and Tennessee Reef), two populations on patch reefs in the inshore channel (East Washerwoman and East Turtle Shoal), and two grass beds located directly offshore in water < 2m in depth (near mile marker 62 on Long Key and behind Keys Marine Lab (KML) on Vaca Key). For broader geographic context we also sampled fishes from two sites further north on the gulf coast of Florida, two sites in the Bahamas, and one site from Belize. Florida and Bahamas specimens were collected in 2005 and 2006, and Belize specimens were collected in 2006. In addition to comparing fore reef and inshore patch reef, we included the grass bed habitat as it experiences even greater seasonal and diurnal fluctuations in temperature than the inshore patch reef and as such provides an additional test of the proposed habitat segregation.

All animal handling procedures were approved by the University of California, Davis Institutional Animal Care and Use Committee. Fish were caught using a combination of hand nets, barrier nets, and otter trawls. Specimens were euthanized using MS-222 dissolved in seawater, and samples were taken from muscle tissue and preserved in 95% ethanol. We extracted DNA using DNeasy™ (Qiagen) columns and PCR amplified a fragment of the mitochondrial cytochrome B gene using the L14768 and H15496 primers from Rocha et al. (2005). PCR products were cleaned using ExoSap-IT (USB Corp.). Purified templates were dye labeled using BigDye (ABI) and sequenced on an ABI 3077 automated DNA Sanger sequencer.

We combined the DNA from our new collections with representative sequences from Rocha et al. (2005) that were available from Genbank (Benson et al. 2013), accession numbers AY823558.1 to AY823569. We aligned sequences using ClustalW (Thompson, Gibson, and Higgins 2002) and inferred a population phylogeny using BEAST v.2.6.3 (Bouckaert et al. 2014). All *cytb* sequences were imported into BEAUTi and partitioned by codon position. All partitions had trees and clocks linked, while site models were allowed to vary. We used ModelTest with “transitionTransversionSplit” (Bouckaert and Drummond 2017) to infer site models. For consistency with Rocha et al. (2005) we also conducted a separate analysis using the TN93 model. All analytical results from the trees inferred with this model were functionally identical to those from the full Bayesian procedure, however, and will not be presented here. We implemented a strict molecular clock and a constant coalescent tree model, as is appropriate for population genetic data when not inferring population size changes (Drummond and Rambaut 2007). BEAST analyses were run twice, with 50,000,000 steps of the Markov chain, sampling every 1000 generations. We constructed a strict consensus tree using the “contree” function in the APE r package, and used it to assign individuals to either “northern” or “southern” haplotypes for visualization and further analysis.

We conducted population genetic analyses using the R packages adegenet and hierfstat (Jombart 2008; Goudet 2005). Because some sites were represented by only a few individuals, we pooled sites by habitat type; “offshore reef”, “inshore reef”, or “inshore grass bed”. To assess whether haplotypes were segregating between different populations we measured pairwise Fst (Nei 1987) between all pairs of habitat types in the Florida Keys. To evaluate the statistical significance of these patterns we compared the observed genetic distance between habitat types to that expected if assortment of haplotypes was random. The expected patterns under this null hypothesis were estimated using a permutation test in which sequences were randomly assigned to habitat types, keeping sample sizes consistent with those from the empirical data. In order to test whether results were robust to our assignment of populations to habitat types we repeated the analyses without pooling sites. Further details of the analysis and all code are provided in the supplemental materials.

Additionally we used a Generalized Mixed-Yule Coalescent model (GMYC) (Pons et al. 2006) for single-locus species delimitation analyses. The GMYC model uses an ultrametric tree to infer a shift between Yule speciation and coalescent processes, using this shift to delimit species. The GMYC method was implemented using the “splits” package in R and the consensus tree inferred using BEAST. We then used a likelihood ratio test to test the hypothesis that more than one species was present in our dataset.

## Results

Phylogenetic and broad-scale biogeographic patterns were concordant with those seen in Rocha et al. (2005), showing a deep divergence between a broadly northern and broadly southern lineage (Figure 1 and Figure 2). We found an approximately equal mix of the two haplotypes in the Florida Keys. In contrast the Bahamas were dominated by the southern haplotype, with only one individual out of the forty having the northern haplotype. We note that this individual was the only Bahamas specimen obtained from Genbank, and that no fine-scale locality information was available for it. Given that the authors providing the original data (Rocha et al. 2005) report the Bahamas as being home to the southern lineage of *H. bivittatus*, however, it is possible that this sequence was misidentified when it was posted to Genbank. Similarly we find that of the two examples from the Virgin Islands in the original study that were available on Genbank, one was from the southern lineage and one was from the northern lineage.

**Figure 1.**
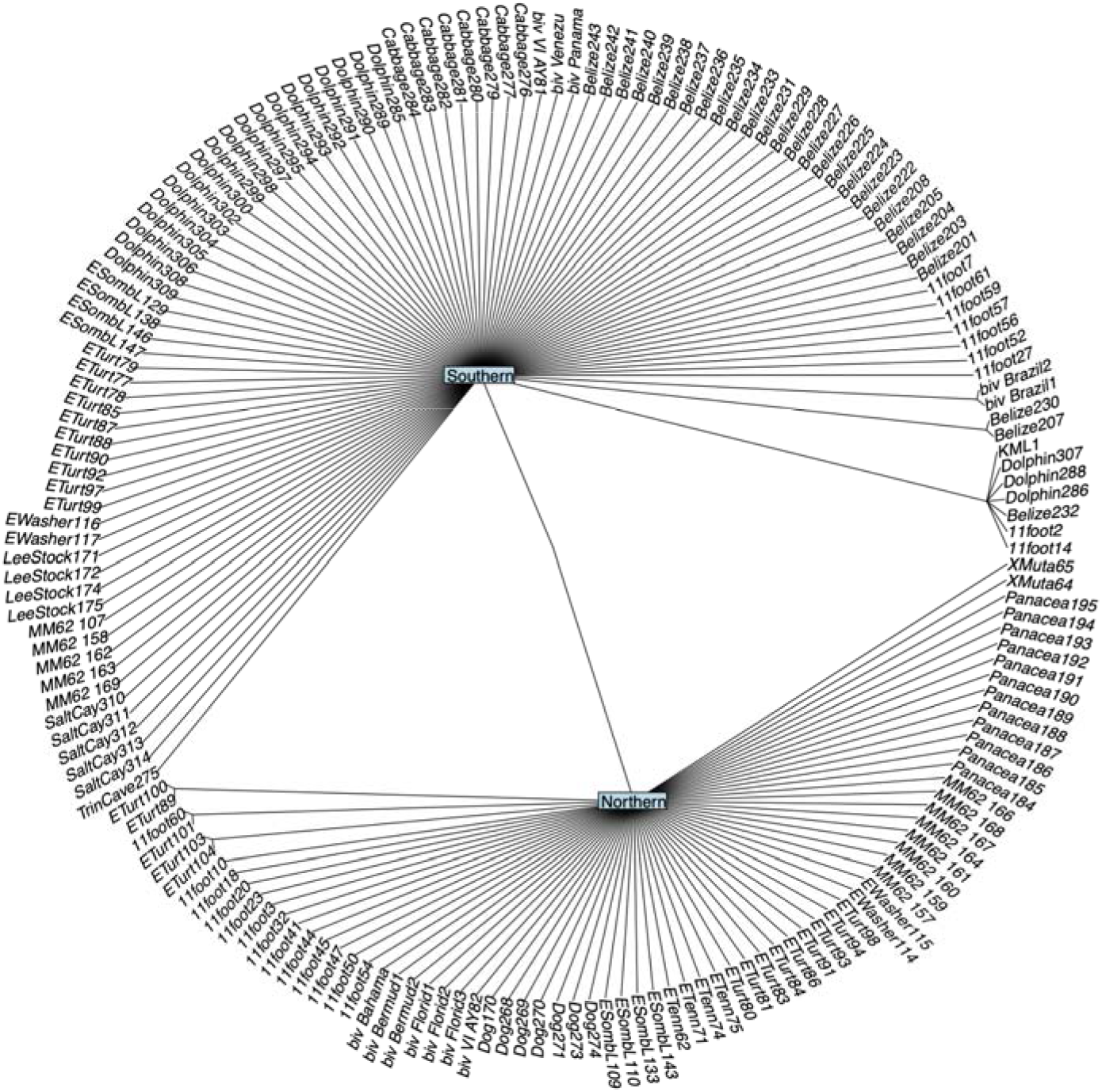
Strict consensus tree showing northern and southern lineages.

**Figure 2.**
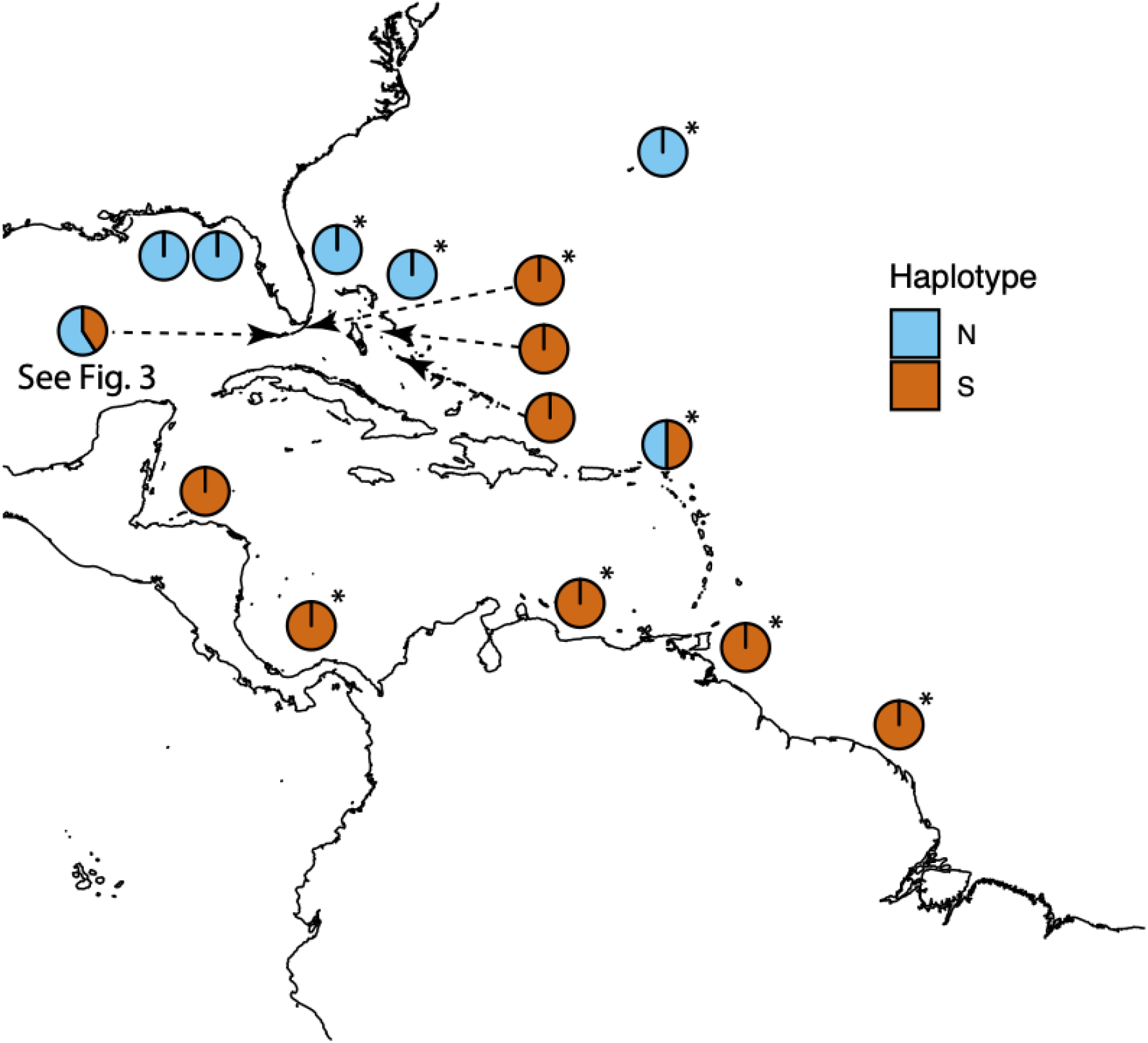
Full study area including Rocha et al. (2005) data from Genbank and new collections. Pie charts indicate relative frequency of haplotypes in different study areas. Pie chart for novel collection data from the Florida Keys is across all newly sampled sites combined; a detailed view of localities within the Keys is given in Figure 3. *Data from Rocha et al. (2005) submissions to Genbank. These were typically one sequence per locality, and do not necessarily represent frequencies reported in the original manuscript.

Our GMYC analysis showed no evidence for more than one species in our dataset (LRT: p = 0.313). Moreover, as shown in Figure 3, the finer-scale analyses of haplotypes in the Florida Keys do not support the hypothesis of habitat segregation presented in Rocha et al. (2005). With higher power to detect differences as a consequence of sampling more individuals in this region (78 specimens vs. 36 specimens), more populations (8 vs. 2) and more diverse habitats (i.e., with the inclusion of fore reef, inshore patch reef, and shallow grass bed populations), we find little evidence for the hypothesis that there are differences in allele frequencies between habitat types or individual populations. Comparison of sites grouped by habitat type showed no significant differences (Figure 4). For the site-level analysis the only statistically significant difference between any pair of populations was between the fore reef site XMuta and the single individual from the KML grass bed site. This result is likely an artifact of permutation tests conducted with a small sample size (4 samples from one population and 1 from the other, see Figure 5). Moreover, in this sole exception, the direction of the difference was opposite to that expected: the specimens sampled from the fore reef were of the northern haplotype, while the lone individual from the shallow grass bed was of the southern haplotype (Figure 3).

**Figure 3.**
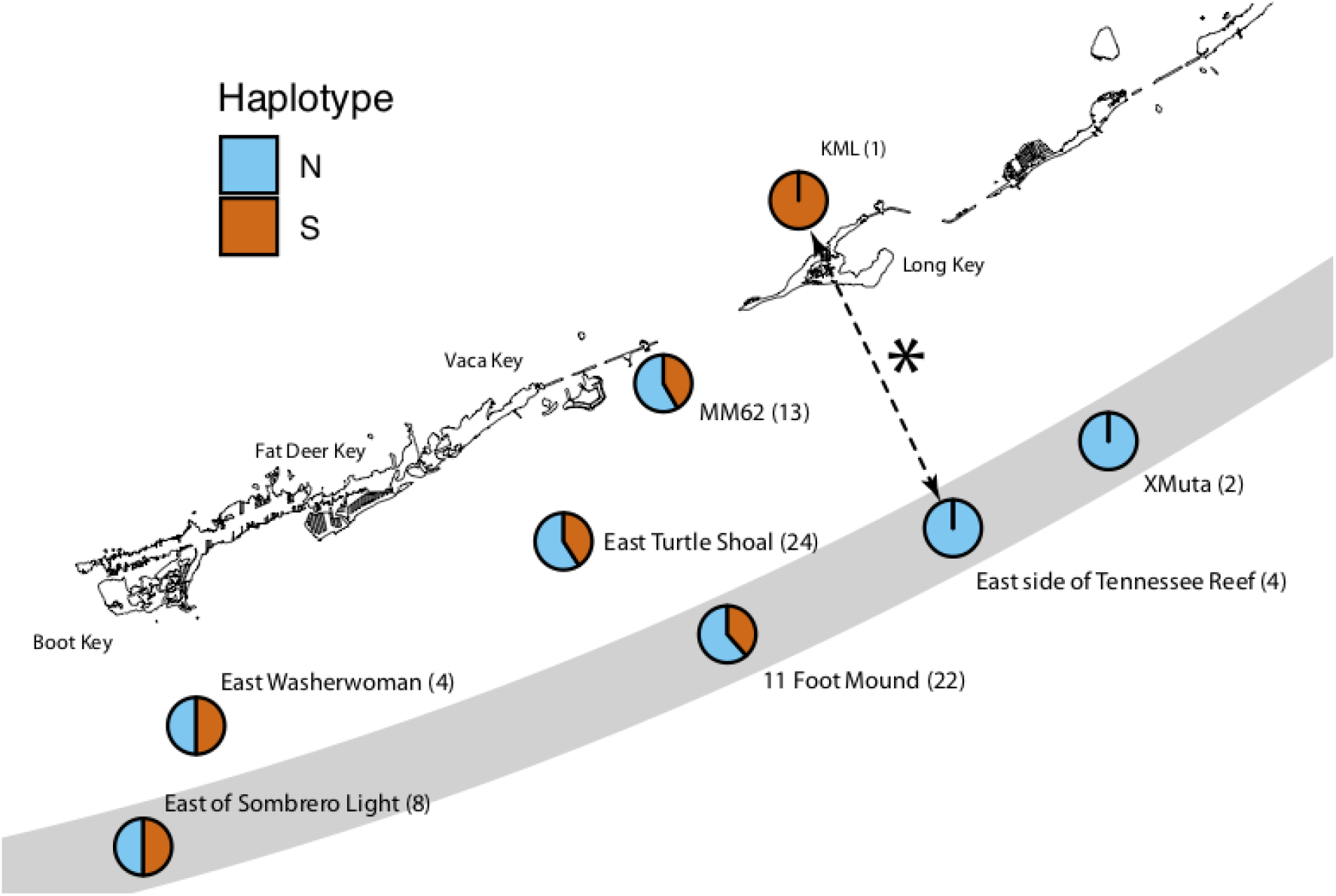
Haplotype frequencies from new sampling localities in the Florida Keys, with sample sizes (in parentheses). Grey bar represents the approximate location of the edge of the continental shelf. Dashed arrow with an asterisk represents the only statistically significant divergence from random assortment of genetic distances (East side of Tennessee Reef vs. KML). This result is opposite to the direction predicted under the Rocha et al. (2005) hypothesis (see main text).

**Figure 4.**
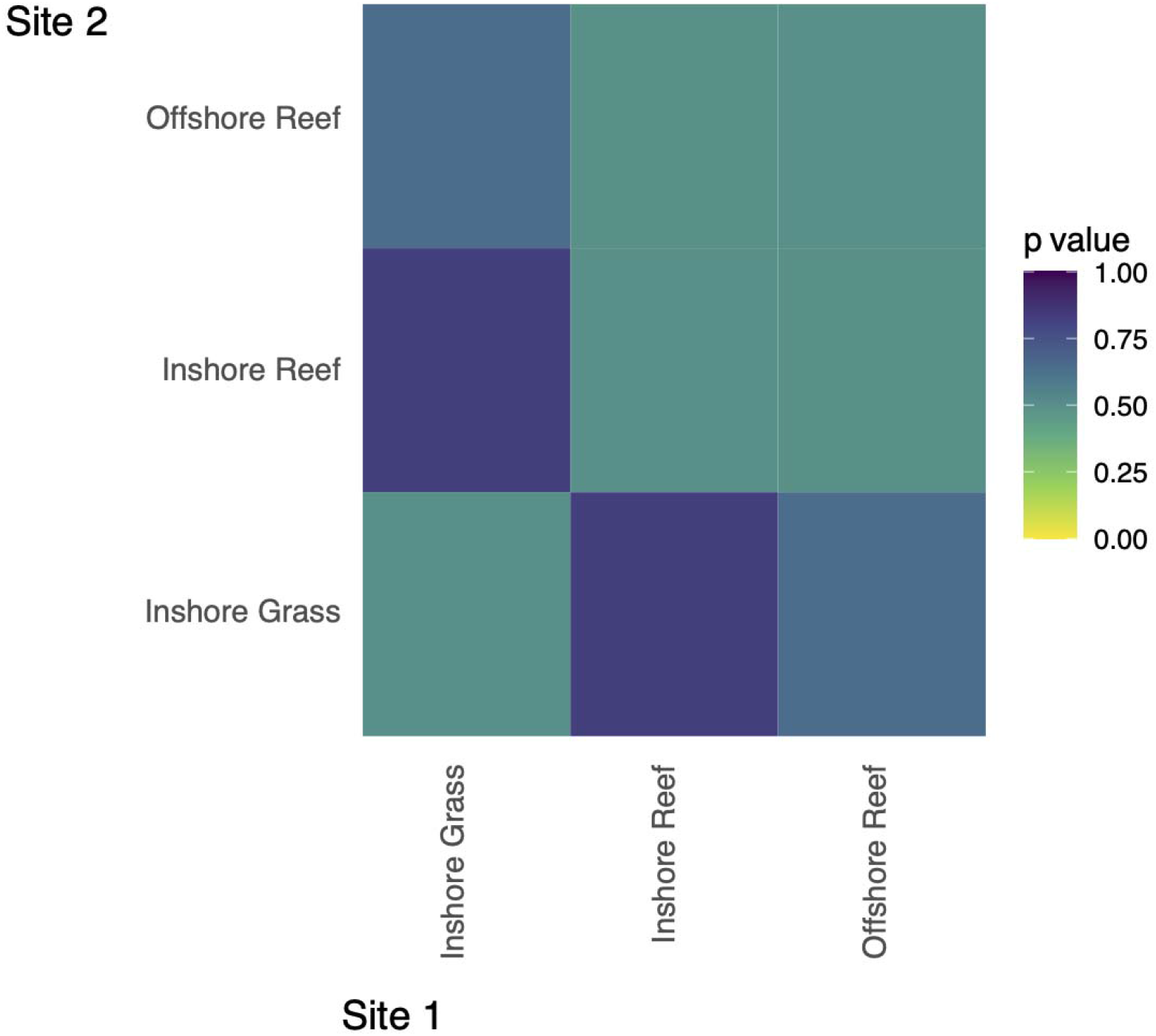
Significance of Fst values from permutation tests, sites grouped by habitat type. Colors represent p values based on 1000 permutations.

**Figure 5.**
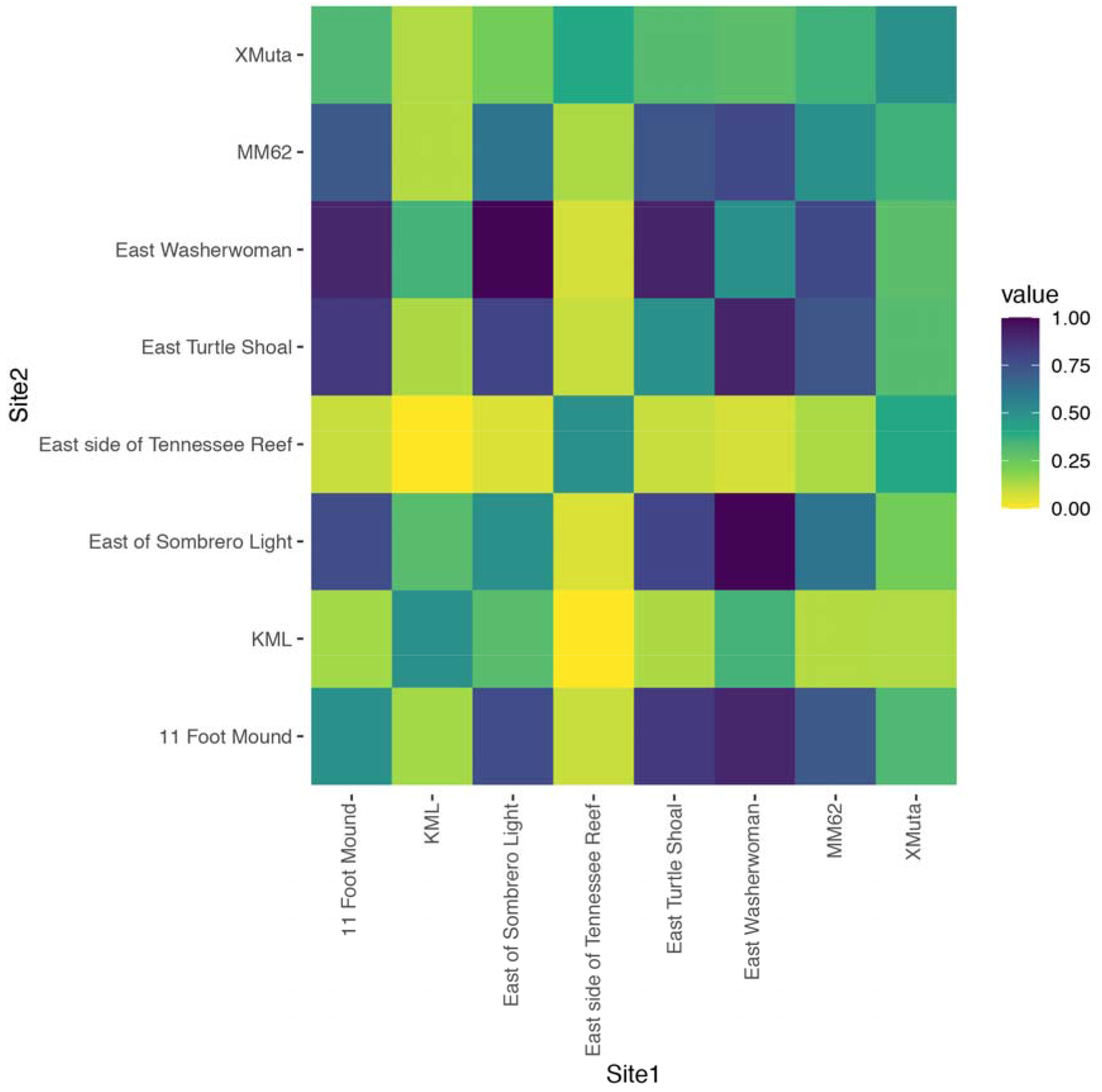
Statistical significance of Fst values from permutation tests, comparing each site individually. Colors represent p values based on 1000 permutations. Only one comparison (East Side of Tennessee Reef vs. KML) was statistically significant at p < .05.

## Discussion

In the current study we attempted to replicate a classic study of parapatric ecological speciation in marine fishes. Our results do not support the hypothesis that the northern and southern lineages of *Halichoeres bivittatus* represent a product of either cryptic or incipient ecological speciation. We find no evidence that these two lineages represent different species. Further we find no evidence for habitat partitioning occuring in the Florida Keys. On the contrary, our study finds that northern and southern lineages are randomly distributed among habitat types and populations in this region. The only site-by-site comparison in the Florida Keys that was significantly different from random assortment was in the opposite direction to that predicted.

Demonstrating speciation with gene flow is notoriously difficult. For these purposes, we find lists of criteria such as those presented by Potkamp and Fransen (2019) to be of particular value; they allow us to quickly quantify the strength of evidence for a given process and adjust our level of confidence accordingly. They suggested six criteria that needed to be addressed:

1. Are populations reproductively isolated?
2. Is there (potential for) disruptive selection?
3. Do populations mate assortatively in sympatry?
4. Is the selected trait linked to the assortment trait?
5. Is there evidence for gene flow between populations at the time of divergence?
6. Do geographic ranges of populations overlap?

Similarly, Nosil (Nosil 2012) established criteria for considering a case of speciation to be “ecological speciation”, which are effectively the same as criteria 1, 2, and 4 above. The case for parapatric speciation and ecological speciation in *Halichoeres bivittatus* so far only consists of direct support for Criterion 6; the presence of both cytochrome B haplotypes in some locations strongly supports the presence of geographically overlapping populations. However, this pattern could also come about if the divergence seen in cytochrome B were entirely due to allopatric divergence followed by secondary contact, or due to neutral processes (Irwin 2002) and as such is not sufficient to support any mechanism of speciation. Consideration of barriers to north/south dispersal that may contribute to allopatric speciation may be particularly relevant, as the contact zone between the two *H. bivittatus* haplotypes in the Florida Keys also mirrors a major faunal break found for many other marine organisms (Robertson and Cramer 2014; Lee and Foighil 2005). Examining the other criteria we find that questions 1 and 3 have not been addressed in any study, while the remainder are supported only by verbal arguments based on the dispersal capability of the group (criterion 5) and the previous finding of habitat assortment in the Keys and Bermuda (criteria 2 and 4).

As we could not replicate the sampling of Rocha et al. (2005) in Bermuda or Key Largo, it is still possible that habitat partitioning is occurring in those localities. In light of our findings and the lack of any demonstration of morphological differentiation or assortative mating between northern and southern lineages, however, we find it difficult to see how such highly localized habitat partitioning could be considered evidence for either ecological speciation or speciation with gene flow in the rest of the Caribbean and Gulf of Mexico. Instead, we caution that differences in haplotype frequencies in these populations could be driven by a number of processes that are not necessarily associated with speciation including lottery recruitment and post-recruitment selection related to local conditions (Grorud-Colvert and Sponaugle 2011; Selkoe et al. 2006; Bernardi et al. 2012; Searcy and Sponaugle 2001). The use of sequences from larvae in Rocha et al. (2005) in some populations makes it particularly difficult to eliminate these processes as alternative explanations for patterns seen in *H. bivittatus*. As such, we would suggest that even those localized results should be viewed with extreme caution until they have been replicated with adult fish over a longer time scale.

There is a growing body of evidence that ecological factors play an important role in in structuring the genetic diversity of marine populations and promoting speciation (Prada and Hellberg 2020; Taylor and Hellberg 2005; Whitney, Donahue, and Karl 2018; Momigliano et al. 2017; Teske et al. 2019; Holt et al. 2020; Bird et al. 2011; Choat et al. 2012; Potkamp and Fransen 2019; Faria, Johannesson, and Stankowski 2021), and failure to replicate one study is not sufficient cause to question the growing consensus that ecological speciation and speciation with gene flow play an important role in generating marine biodiversity. Likewise, there is an abundance of evidence that allopatry has also promoted speciation in marine settings (Wainwright et al. 2018; Ekimova et al. 2019; Chenuil et al. 2018; Holt et al. 2020; Laakkonen et al. 2021). We should not be surprised that in such a species-rich and unique environment there is evidence for a variety of speciation mechanisms. The question is when should we conclude that the weight of evidence supports a given scenario. While many in the field might suggest that our default position should be one of assuming allopatric speciation until proven otherwise, we are less confident that this is the appropriate stance to take for marine environments. Rather we suggest that when we don’t know the answer to four of the six criteria for demonstrating speciation with gene flow, or for any of the three criteria for demonstrating ecological speciation, the most appropriate position is to simply acknowledge that we have insufficient evidence to argue for any mechanism of speciation in this system. “We don’t know” is a deeply unsatisfying answer, but it’s the only one that accurately reflects the currently available evidence.

## Supporting information

Code and visualization for analyses

## Acknowledgements

The authors would like to thank the generous assistance of the Keys Marine Lab, FSU Coastal and Marine Laboratory, Dolphin Encounters, and Gulf Specimen Marine Laboratory. We would also like to thank Lisa Tipsword, Lonnie Anderson, and Colleen Young for assistance in the field, as well as the help of Kyle Miller from the Florida Fish and Wildlife Conservation Commission and Roland Albury in the Bahamas Department of Marine Resources (Fisheries). The authors are also grateful to Luiz Rocha, who provided very helpful feedback on an earlier version of the manuscript.

## Author Contributions

The study was conceived and designed by PCW and DLW. Fieldwork was conducted by all authors. DNA extraction and sequencing was performed by MCB and DLW. DLW and RIE performed data analyses. All authors contributed to writing the manuscript.

