## Supplementary material for "Reevaluating claims of ecological speciation in *Halichoeres bivittatus*": Code and visualization for analyses: Hal-biv-biogeography.html

Halichoeres Bivitattus Biogeography


### Halichoeres Bivitattus Biogeography

### Setup

```
library(babette)
library(mcbette)
library(beastier)
library(ape)
library(knitr)
library(phytools)
library(geiger)
library(DT)
library(sf)
library(leaflet)
library(tidyr)
library(plyr)
library(dplyr)
library(leaflet.minicharts)
library(ggplot2)
library(reshape2)
library(adegenet)
library(hierfstat)

setwd("~/Dropbox/Ongoing Projects/Hal Biv biogeography/")
```

### Introduction

In this analysis we’re going to take an alignment of cytochrome B from a bunch of *Halichoeres bivittatus* specimens, build a tree, and ask whether there’s any indication that haplotypes are segregated by habitat in the Florida Keys. The first part of this analysis is very heavily based on this ROpenSci post by Richèl J.C. Bilderbeek

### Alignment

First we’ll load in our alignment and visualize it to make sure there aren’t any obvious problems.

```
save_nexus_as_fasta("Halicheoeres NEW cytb_no_Out.nex",
                    "halbiv.fasta")

par0 <- par(mar = c(3, 7, 3, 1))
dna_sequences <- read.FASTA("halbiv.fasta")
image.DNAbin(dna_sequences, mar = c(3, 7, 3, 1))
```

### Building the tree

*Note* We’re going to need to change the inference model etc. to match what Ron did; this is my older version listed here.

```
inf.model <- create_inference_model(
  site_model = create_tn93_site_model()
)
trees <- bbt_run_from_model("halbiv.fasta",
                            inference_model = inf.model)
save(trees, file = "halbiv-beast.Rda")
contree <- consensus(trees$halbiv_trees[1001:10001])
save(contree, file = "contree.Rda")
```

```
trees <- read.nexus("./HalichoeresNEWcytb_no_OutStrictCLKbMTnoCalibration.trees")
save(trees, file = "halbiv-beast.Rda")
contree <- consensus(trees[seq(1001, 50001, by = 10)])
save(contree, file = "contree.Rda")
```

#### Densitree plot

```
load("halbiv-beast.Rda")
load("contree.Rda")

plot_densitree(
  trees[seq(1001, 50001, by = 1000)],
  alpha = 0.1
)
```

```
pdf("densitree.pdf", height = 34)
plot_densitree(
  trees[seq(1001, 50001, by = 1000)],
  alpha = 0.1
)
dev.off()
```

```
## quartz_off_screen 
##                 2
```

As expected based on previous analyses and the Rocha et al. results, there’s a very deep split here that is very highly supported. There’s little resolution at shallow nodes, but the north/south split is very strong. The lineage at the top constitutes the “northern” haplotype, and includes the specimens from northern Florida (Dog Island, Panacea, St. Petersburg) as well as some of the specimens from the Bahamas and Bermuda. It also includes about half of the specimens from the Keys. The southern split, which also contains about half of the Keys specimens, also contains the specimens from Venezuela, the Virgin Islands, Panama, Brazil, some of the Bahamas specimens, and Belize.

Thus at the broad geographic scale the overall story here is the same as in Rocha et al.; a very deep split between a northern and southern haplotype, with a mix of the two at a suture zone in the Keys and the Bahamas. Rocha et al. said there’s also a mix in Bermuda, but we don’t have the data to address that issue here.

#### Strict consensus tree

In order to condense this down to N/S haplotype assignments, we’ll take a strict consensus tree from that MCMC run.

```
plot(contree, type = "radial", lab4ut = "axial")
nodelabels(c("Southern", "Northern"), node = c(179, 183))
```

```
pdf("contree.pdf", width = 15, height = 15)
plot(contree, type = "radial", lab4ut = "axial")
nodelabels(c("Southern", "Northern"), node = c(179, 183))
dev.off()
```

```
## quartz_off_screen 
##                 2
```

From examining the tree we can see that the southern haplotype is represented by the clade descending from node 179 and the northern haplotype is node 184.

#### Extracting clade membership

```
habitat.df <- read.csv("habitat by specimen.csv")
habitat.df$haplotype <- rep(NA, nrow(habitat.df))
habitat.df[habitat.df$Specimen %in% tips(contree, 179),"haplotype"] <- "S"
habitat.df[habitat.df$Specimen %in% tips(contree, 183),"haplotype"] <- "N"

datatable(habitat.df, extensions = 'Buttons',
          options = list(dom = 'Blfrtip',
                         buttons = c('copy', 'csv', 
                                     'excel', 'pdf', 'print'),
                         columnDefs = list(list(className = 'dt-center',
                                                targets = 0:ncol(habitat.df))),
                         lengthMenu = list(c(10,25,50,-1),
                                           c(10,25,50,"All"))))
```

#### Data aggregation

```
localities <- read_sf("./doc.kml") %>%
  as_Spatial() %>%
  as.data.frame() %>%
  select(Name, coords.x1, coords.x2) %>%
  filter(!Name == "East East Turtle Shoal")

colnames(localities) <- c("Site", "Longitude", "Latitude")

hapcount.df <- habitat.df %>%
  na.omit() %>%
  dplyr::count(Site, Habitat.type, Location, haplotype) %>%
  spread(key = haplotype, value = n, fill = 0)

localities <- join(localities, hapcount.df, by = "Site")
```

#### Plotting sampling locations

```
datatable(localities, extensions = 'Buttons',
          options = list(dom = 'Blfrtip',
                         buttons = c('copy', 'csv', 
                                     'excel', 'pdf', 'print'),
                         columnDefs = list(list(className = 'dt-center',
                                                targets = 0:ncol(localities))),
                         lengthMenu = list(c(10,25,50,-1),
                                           c(10,25,50,"All")))) %>%
  formatRound(columns=c("Longitude", "Latitude"), digits=3)
```

#### Haplotype frequencies by sampling site

Since we don’t have the non-Florida samples in the KMZ file, I’m going to add them manually for plotting purposes. In the sampling file we have a bunch of samples labled “Dolphin Encounters” and some others labels “Salt Cay”. The “Dolphin Encounters” samples were collected by DW and RE, and I think the “Salt Cay” ones were collected by PW. Dolphin Encounters is actually on Salt Cay, though, so we’re going to merge those together for plotting purposes.

Note that some of these are based on samples posted to Genbank by Rocha et al., and those only have country names associated with them, not latitudes and longitudes. Placement of those pie charts is therefore approximate.

```
colors <- c("#f5dc7a", "#a532ed")

all.localities <- read.csv("all.localities.csv")

leaflet(all.localities) %>%
  addProviderTiles(provider = "GeoportailFrance.orthos") %>%
  addCircleMarkers(label = ~ Site, opacity = 0, fillOpacity = 0,
                   radius = 20) %>%
  addMinicharts(all.localities$Longitude, all.localities$Latitude,
                type = "pie",
                chartdata = all.localities[,c("N", "S")],
                colorPalette = colors)
```

### Habitat segregation in the Florida Keys

#### Haplotpye frequencies, Keys only

```
colors <- c("#f5dc7a", "#a532ed")

leaflet(localities) %>%
  addProviderTiles(provider = "GeoportailFrance.orthos") %>%
  addCircleMarkers(label = ~ Site, opacity = 0, fillOpacity = 0,
                   radius = 20) %>%
  addMinicharts(localities$Longitude, localities$Latitude,
                type = "pie",
                chartdata = localities[,c("N", "S")],
                colorPalette = colors)
```

Here we see that both haplotypes are found at somehwat high frequencies at most sites. The two exceptions (KML, X Muta, and the east side of Tennessee Reef) are the two with the smallest sample sizes (1, 2, and 4 samples, respectively). As a result it’s probably best not to read too much into those sites.

This next bit of code is just re-making the pie charts so we can assemble a vector map in Illustrator.

```
all.localities.long <- melt(all.localities, 
                            id.vars = c("Site", "Longitude", "Latitude", "Habitat.type", "Location"), 
                            variable.name = "Haplotype")

for(i in unique(all.localities.long$Site)){
  this.df <- all.localities.long %>%
    filter(Site == i)
  
  pdf(file = paste0(i, " pie chart.pdf"))
  print(ggplot(this.df, aes(x="", y=value, fill=Haplotype)) +
    geom_bar(stat="identity", width=2, color = "black") +
    coord_polar("y", start=0) + theme_void() + 
    scale_fill_manual(values = c("#404040", "#D0D0D0")))
  dev.off()
}
```

#### By habitat type

##### Haplotype frequencies by habitat type

```
localities %>% 
  pivot_longer(c("N", "S"), names_to = "haplotype", values_to = "count") %>%
  ggplot(aes(fill=haplotype, y=count, x=Habitat.type)) + 
  geom_bar(position="fill", stat="identity")
```

```
pdf("freq.by.type.pdf")
localities %>% 
  pivot_longer(c("N", "S"), names_to = "haplotype", values_to = "count") %>%
  ggplot(aes(fill=haplotype, y=count, x=Habitat.type)) + 
  geom_bar(position="fill", stat="identity")
dev.off()
```

```
## quartz_off_screen 
##                 2
```

In this plot we’re comparing haplotype frequencies for shallow (< 2m) grass bed communities to those for reef communities, combining both patch reef and fore reef. Again we see very similar frequencies.

##### Genetic distance between habitat types

We’re going to measure Fst between all habitat types in the Florida Keys, and then compare those values to the distribution of pairwise Fsts we get from permutation tests. For these permutations, we simply randomly permute the population vector and then measure Fsts again, giving us a distribution of expected pairwise Fsts under the hypothesis that there is no habitat segregation. We’re doing 1000 permutations.

```
fl.sequences <- subset(dna_sequences, names(dna_sequences) %in%
                         habitat.df[complete.cases(habitat.df),"Specimen"])
halbiv.genind <- DNAbin2genind(fl.sequences)
fl.specimens <- rownames(halbiv.genind$tab)
fl.habitat.df <- habitat.df[habitat.df$Specimen %in% fl.specimens,]
rownames(fl.habitat.df) <- fl.habitat.df$Specimen
fl.habitat.df$full.habitat <- paste(fl.habitat.df$Location,
                                    fl.habitat.df$Habitat.type)

halbiv.genind$pop <- as.factor(fl.habitat.df[fl.specimens, "full.habitat"])

fst.dist <- as.matrix(genet.dist(halbiv.genind, method="Nei87"))

if(file.exists("habitat.fst.rank.csv")){
  habitat.rank.df <- read.csv("habitat.fst.rank.csv", row.names = 1)
  colnames(habitat.rank.df) <- rownames(habitat.rank.df)
} else {
  
  sims <- list()
  nsims <- 1000
  
  for(i in 1:nsims){
    this.genind <- halbiv.genind
    this.genind$pop <- sample(this.genind$pop)
    sims[[i]] <- as.matrix(genet.dist(this.genind, method="Nei87"))
  }
  
  # Step through each row/column combo and figure out the 
  # ranking of our empirical result in the simulations.
  
  habitat.rank.df <- as.data.frame(fst.dist)
  habitat.rank.df[,] <- NA
  
  for(i in 1:nrow(fst.dist)){
    for(j in 1:ncol(fst.dist)){
      habitat.rank.df[i,j] <- rank(c(fst.dist[i,j], sapply(sims, function(x) x[i,j])))[1]
    }
  }
  
  habitat.rank.df <- 1 - habitat.rank.df/(nsims + 1)
  
  write.csv(habitat.rank.df, file = "habitat.fst.rank.csv")
}

datatable(habitat.rank.df, extensions = 'Buttons',
          options = list(dom = 'Blfrtip',
                         buttons = c('copy', 'csv', 
                                     'excel', 'pdf', 'print'),
                         columnDefs = list(list(className = 'dt-center',
                                                targets = 0:ncol(habitat.rank.df))),
                         lengthMenu = list(c(10,25,50,-1),
                                           c(10,25,50,"All")))) %>%
  formatRound(columns=colnames(habitat.rank.df), digits=3)
```

```
melted.cor.mat <- reshape2::melt(as.matrix(habitat.rank.df))
colnames(melted.cor.mat) <- c("Site1", "Site2", "value")

cor.heatmap <- ggplot(data = melted.cor.mat, aes_string(x="Site1", y="Site2", fill="value")) +
  geom_tile() + scale_fill_viridis_c(direction = -1, limits = c(0,1)) +
  theme(axis.text.x = element_text(angle = 90, hjust = 1)) + 
  theme(panel.grid.major = element_blank(), panel.grid.minor = element_blank(),
        panel.background = element_blank())
cor.heatmap
```

```
pdf("fst.by.type.pdf")
cor.heatmap
dev.off()
```

```
## quartz_off_screen 
##                 2
```

#### By site

##### Haplotype frequencies by site category (onshore vs. offshore)

```
localities %>% 
  pivot_longer(c("N", "S"), names_to = "haplotype", values_to = "count") %>%
  ggplot(aes(fill=haplotype, y=count, x=Location)) + 
  geom_bar(position="fill", stat="identity")
```

```
pdf("hap.freqs.by.site")
localities %>% 
  pivot_longer(c("N", "S"), names_to = "haplotype", values_to = "count") %>%
  ggplot(aes(fill=haplotype, y=count, x=Location)) + 
  geom_bar(position="fill", stat="identity")
dev.off()
```

```
## quartz_off_screen 
##                 2
```

Here we see that the haplotype frequencies in onshore and offshore sites are not very different from one another, and to the extent that they do differ it is in the opposite direction predicted by the Rocha et al. (2015) hypothesis; we find slightly higher prevalence of the southern haplotype at the inshore patch reefs and grass bed sites, which are expected to experience greater fluctuations in temperature.

##### Genetic distance between sites

We’re going to measure Fst between all sites in the Florida Keys, and then compare those values to the distribution of pairwise Fsts we get from permutation tests. For these permutations, we simply randomly permute the population vector and then measure Fsts again, giving us a distribution of expected pairwise Fsts under the hypothesis that there is no habitat segregation. We’re doing 1000 permutations.

```
halbiv.genind <- DNAbin2genind(fl.sequences)
fl.specimens <- rownames(halbiv.genind$tab)
fl.habitat.df <- habitat.df[habitat.df$Specimen %in% fl.specimens,]
rownames(fl.habitat.df) <- fl.habitat.df$Specimen

halbiv.genind$pop <- as.factor(fl.habitat.df[fl.specimens, "Site"])

fst.dist <- as.matrix(genet.dist(halbiv.genind, method="Nei87"))

if(file.exists("site.fst.rank.csv")){
  site.rank.df <- read.csv("site.fst.rank.csv", row.names = 1)
  colnames(site.rank.df) <- rownames(site.rank.df)
} else {
  
  sims <- list()
  nsims <- 1000
  
  for(i in 1:nsims){
    this.genind <- halbiv.genind
    this.genind$pop <- sample(this.genind$pop)
    sims[[i]] <- as.matrix(genet.dist(this.genind, method="Nei87"))
  }
  
  # Step through each row/column combo and figure out the 
  # ranking of our empirical result in the simulations.
  
  site.rank.df <- as.data.frame(fst.dist)
  site.rank.df[,] <- NA
  
  for(i in 1:nrow(fst.dist)){
    for(j in 1:ncol(fst.dist)){
      site.rank.df[i,j] <- rank(c(fst.dist[i,j], sapply(sims, function(x) x[i,j])))[1]
    }
  }
  
  site.rank.df <- 1 - site.rank.df/(nsims + 1)
  
  write.csv(site.rank.df, file = "site.fst.rank.csv")
}

datatable(site.rank.df, extensions = 'Buttons',
          options = list(dom = 'Blfrtip',
                         buttons = c('copy', 'csv', 
                                     'excel', 'pdf', 'print'),
                         columnDefs = list(list(className = 'dt-center',
                                                targets = 0:ncol(site.rank.df))),
                         lengthMenu = list(c(10,25,50,-1),
                                           c(10,25,50,"All")))) %>%
  formatRound(columns=colnames(site.rank.df), digits=3)
```

```
melted.cor.mat <- reshape2::melt(as.matrix(site.rank.df))
colnames(melted.cor.mat) <- c("Site1", "Site2", "value")

cor.heatmap <- ggplot(data = melted.cor.mat, aes_string(x="Site1", y="Site2", fill="value")) +
  geom_tile() + scale_fill_viridis_c(direction = -1, limits = c(0,1)) +
  theme(axis.text.x = element_text(angle = 90, hjust = 1)) + 
  theme(panel.grid.major = element_blank(), panel.grid.minor = element_blank(),
        panel.background = element_blank())
cor.heatmap
```

```
pdf("fst.by.site.pdf")
cor.heatmap
dev.off()
```

```
## quartz_off_screen 
##                 2
```

Just for simplicity’s sake I’m going to set all p > .05 to 1 and re-plot that heatmap to highlight which comparisons are significant.

```
melted.cor.mat[which(melted.cor.mat$value > .05), "value"] <- 1
cor.heatmap <- ggplot(data = melted.cor.mat, aes_string(x="Site1", y="Site2", fill="value")) +
  geom_tile() + scale_fill_viridis_c(direction = -1, limits = c(0,1)) +
  theme(axis.text.x = element_text(angle = 90, hjust = 1)) + 
  theme(panel.grid.major = element_blank(), panel.grid.minor = element_blank(),
        panel.background = element_blank())
cor.heatmap
```

```
pdf("fst.by.site.sigonly.pdf")
cor.heatmap
dev.off()
```

```
## quartz_off_screen 
##                 2
```

So the only comparison that is statistically significant (without Bonferroni correction) is the comparison between East Tennessee Reef and the grass bed behind KML. However the sample sizes are miniscule there (4 vs. 1), and the direction of difference seen is exactly opposite to that predicted by the Rocha et al. result, with the northern haplotype dominating out on the dropoff and the KML specimen being the southern haplotype.
